# NSD1 governs H3K36me2-mediated DNA methylation and drives differentiation of human iPSCs by regulating ‘HIDEN’ lncRNA expression

**DOI:** 10.1101/2025.06.16.659867

**Authors:** Anna Hansknecht, Martina Mankarious, Monica Varona Baranda, Esra Dursun Torlak, Wolfgang Wagner, Kira Zeevaert, Deepika Puri

## Abstract

Epigenetic regulatory mechanisms, which include histone modifications and DNA methylation, play a central role in development and aging. Dimethylation of H3K36, deposited mainly by the histone methyltransferase NSD1, occurs predominantly in intergenic regions and recruits the DNA methyltransferase DNMT3A to facilitate DNA methylation. Haploinsufficiency of NSD1 results in the overgrowth disorder Sotos syndrome *in vivo*, which is associated with aberrant DNA methylation signatures and an enhanced epigenetic age. To understand the mechanisms by which NSD1 may regulate development, differentiation and diseases, we generated human iPSC lines deficient in functional NSD1 (NSD1-KO). NSD1-KO cells exhibit a substrate-specific decrease in proliferation, reduced H3K36me2 levels and extensive DNA hypomethylation. Notably, the loss of functional NSD1 altered the differentiation potential of iPSCs, with aberrant endodermal and mesodermal lineage commitment. Further analysis revealed that NSD1 might drive endodermal differentiation by regulating the expression of an endodermal lncRNA ‘HIDEN’ by mechanisms independent of the regulation of DNA methylation. Our NSD1-KO iPSC lines partially recapitulate the DNA methylation defects associated with NSD1-related disorders. Additionally, we uncovered a novel mechanism by which NSD1 may regulate endodermal differentiation of human iPSCs.

## Introduction

Epigenetic modifications are crucial in determining the activity and accessibility of genes and regulatory elements without altering the underlying DNA sequence [1]. Interactions between multiple epigenetic features, such as DNA methylation and histone modifications, play a central role during cellular development and maturation [2–4], and aberrations in these pathways contribute to the pathogenesis of various diseases, including cancer [5–7]. Therefore, understanding the epigenetic network is critical for elucidating the mechanisms governing differentiation, aging, and the development of pathogenic transcriptional activity.

In mammals, CG dinucleotides (CpG sites) are methylated in a regulated manner during cellular development and aging [8–11]. Recent studies have implicated histone modifications such as Histone H3 lysine 36 (H3K36) methylation in the recruitment of DNMT proteins and consequently the establishment of DNA methylation [12]. Dimethylation of H3K36 (H3K36me2) predominantly colocalizes with DNMT3A, especially in intergenic regions, playing a key role in the establishment of the DNA methylation pattern [12,13]. H3K36me2 is deposited by several histone methyltransferases (HMTs), however, the Nuclear Receptor Binding SET Domain Protein 1 (NSD1) is one of the major enzymes establishing this histone mark in humans [12]. Underlining the role of NSD1 for early differentiation processes, homozygous NSD1-deficient mouse embryos display an impaired mesendoderm formation and fail to complete gastrulation [14]. Furthermore, studies in mouse embryonic stem cells implicate NSD1 function in stem cell differentiation processes [15,16]. In humans, NSD1 dysfunction is linked to diseases such as Sotos syndrome, in which over 90% of the patients have abnormalities in NSD1 [17,18]. Sotos syndrome is a childhood overgrowth syndrome characterized by distinctive facial features, physical overgrowth, advanced bone age, and learning disabilities [19,20]. Genome-wide DNA methylation analysis in Sotos syndrome patients revealed a specific signature that distinguishes pathological NSD1 mutations from benign or control patterns, global DNA hypomethylation, especially in intergenic regions, and an acceleration of epigenetic age [12,21,22].

NSD1-mediated H3K36me2 has been reported to restrict PRC2 activity and prevent aberrant polycomb-mediated gene silencing and early differentiation of stem cells through uncontrolled deposition of H3K27me3 [23]. Furthermore, reports indicate that NSD1 can bind to methylated H3K4 and H3K9, potentially impacting gene expression [24]. NSD1 also binds to enhancers and modulates the H4K27Ac levels, hence regulating enhancer activity [15,16,25]. These reports and more indicate that NSD1 is part of a complex interplay of epigenetic regulation that may play critical roles in gene expression and cellular function.

To better understand the interplay between NSD1-mediated H3K36me2 and DNA methylation and its impact on development and differentiation, we modified NSD1 in human induced pluripotent stem cells (iPSCs) using CRISPR/Cas9 gene editing and systematically reduced H3K36me2. Human iPSCs have the capacity for cellular differentiation into the three germ layers and all derived cell lines, such as mesenchymal stromal cells, which make them a valuable tool for disease modeling, analysis of early development and senescence-associated epigenetic changes [26]. NSD1 modified cell lines retained pluripotency markers, whereas differentiation potential towards endodermal and mesodermal lineages was impaired. We found a significant downregulation of the Human IMP1-Associated “Desert” Definitive Endoderm lncRNA (*HIDEN*), which plays a major role in endodermal differentiation of human pluripotent cells mediated by the WNT signaling pathway [27]. Furthermore, extensive DNA hypomethylation at intergenic regions showed overlapping patterns with Sotos syndrome patients. Overall, our results provide valuable insights into NSD1-mediated H3K36 dimethylation as a regulator for DNA methylation and differentiation potential of human iPSCs.

## Results

### Genetic modification of *NSD1* induces loss of H3K36 dimethylation in human iPSCs

Exon three of the human *NSD1* gene encodes for the N-terminal PWWP domain of canonical NSD1 and was targeted in human iPSCs for CRISPR/Cas9 gene editing (Supplementary Figure 1A). Three iPSC lines (NSD1-1, NSD1-2, NSD1-3) were generated, and frameshift mutations were confirmed via Sanger Sequencing (Supplementary Figure 1A). While we were still able to detect NSD1 by immunophenotypic analysis (Supplementary Figure 1B), the generated iPSC lines revealed a loss of NSD1 function, having reduced H3K36me2 in Western Blot and immunophenotypic analysis (Figure 1A, Supplementary Figure 1C). ChIP-Seq analysis for H3K36me2 also showed significantly reduced enrichment in at least two of the three clones (clone 2 and 3) (Figure 1B, Supplementary Figure 1D), which was especially pronounced in enhancer regions (Figure 1C). Further annotation of H3K36me2 regions lost in NSD1-KO cells revealed that more than 50% mapped to intronic and intergenic regions (Supplementary Figure 1E). Despite the significant loss in H3K36me2, NSD1-KO iPSCs retain pluripotency markers (Figure 1D), which was confirmed at the DNA methylation level by a positive Epi-Pluri-Score, which classifies pluripotent and somatic cells [28]. Notably, NSD1-KO iPSCs demonstrated significantly reduced growth compared to WT iPSCs on the common maintenance substrate Vitronectin (VTN) (Figure 1E). We investigated whether alternative substrates could be used for culture and observed that growth was comparable to WT iPSCs when cultivating NSD1-KO iPSCs on Biolaminin 521 LN (LMN) (Supplementary Figure 1F), and the cells were grown on LMN for further experiments. Taken together, our NDS1-KO iPSCs show reduced H3K36me2 enrichment in intergenic regions and exhibit significant growth defects.

**Figure 1:**
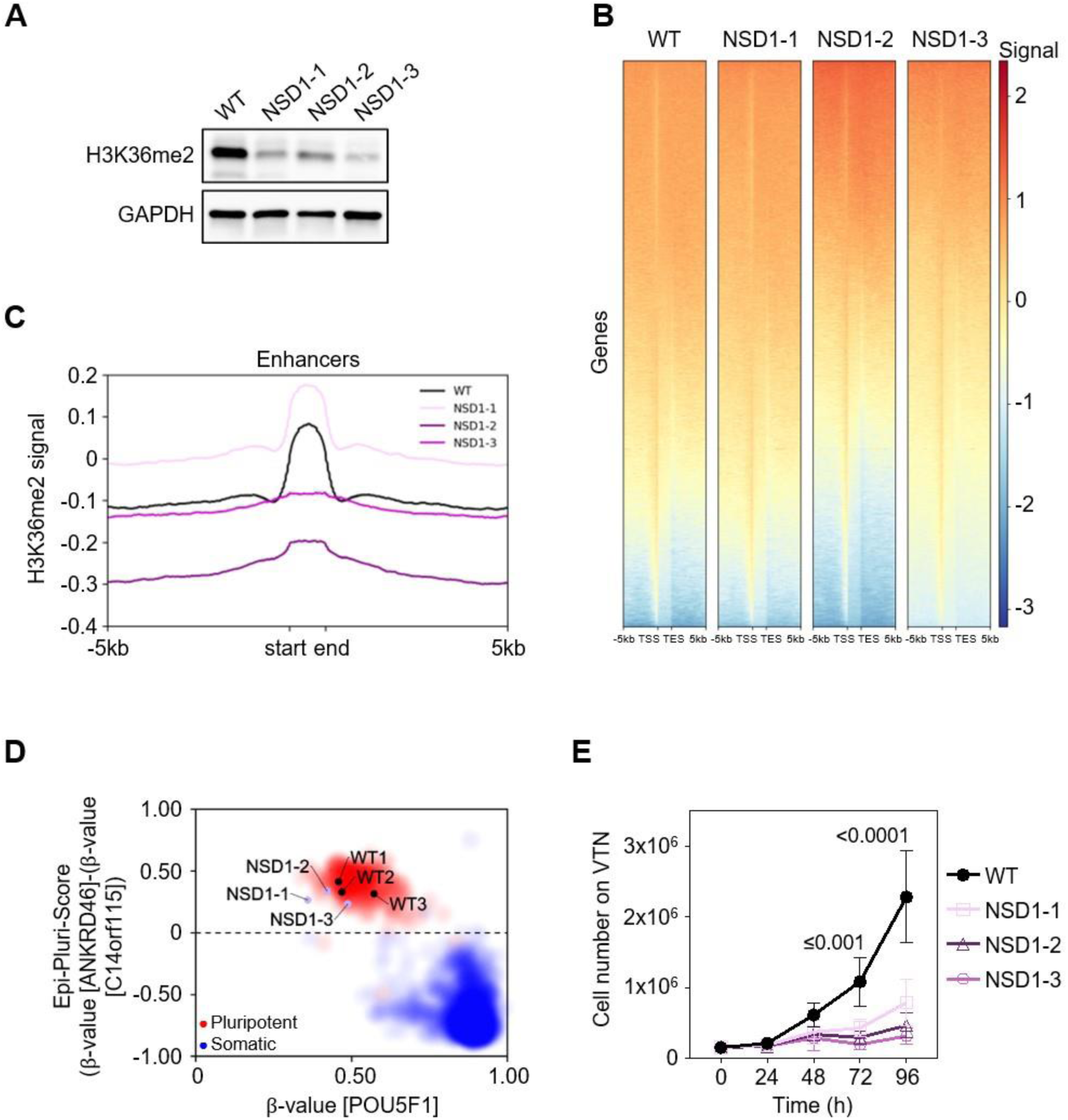
NSD1-KO iPSCs show reduced H3K36me2. **(A)** Western Blot analysis of WT and the three NSD1-KO iPSC lines using antibodies targeting H3K36me2 and the housekeeping protein GAPDH. **(B)** Heatmap of H3K36me2 ChIP-seq signal showing transcription start site (TSS) and transcription end site (TES) ±5 kbp for WT and NSD1-KO clones. **(C)** Line plot showing H3K36me2 signal intensity over enhancer regions ±5 kbp for WT and NSD1-KO clones. **(D)** Epi-Pluri-Score of WT (n=3) and NSD1-KO iPSCs. This analysis assesses DNA methylation at three specific CpG sites, including one within the pluripotency gene *POU5F1* (also known as OCT4). Methylation differences in *ANKRD46* and *C14orf115* are combined to calculate the Epi-Pluri-Score. Background dots represent DNA methylation profiles from 264 pluripotent (red) and 1951 non-pluripotent (blue) cell samples [28]. **(E)** Growth curves of WT and the three NSD1-KO iPSC lines on Vitronectin for 96h. Each time point was measured in technical triplicate. Significance was determined by a Two-way ANOVA test, and p-values are depicted.

### NSD1-KO induces global DNA hypomethylation, similar to Sotos syndrome

Previous reports have shown that loss of NSD1-mediated H3K36me2 results in DNA hypomethylation, specifically at intergenic sites, and causes DNMT3A to relocalize to the H3K36me3 mark within active gene bodies in mouse embryonic stem cells [12]. To further elucidate the effect of reduced NSD1-mediated H3K36me2 in human iPSCs, we performed Illumina Beadchip-based DNA methylation analysis. WT and NSD1-KO cell lines clustered apart from each other in the MDS plot (Supplementary Figure 2A). We observed significant (mean beta value difference ≥ 0.2 and adjusted p-values ≤ 0.05 as significant) DNA hypomethylation at 14,043 CpG sites (Figure 2A). The hypomethylated CpGs mapped largely to intergenic regions (Figure 2B) and revealed associations with Gene Ontology (GO) terms, such as plasma membrane, monoatomic ion transport, or transporter activity (Supplementary Figure 2B). In contrast, 607 CpGs were hypermethylated in NSD1-KO cells (Figure 2A) and mapped to gene body and 3′-UTR regions with less significant GO terms (Figure 2B, Supplementary Figure 2C). Comparison with ChIP-sequencing data indicated that 34.9% of hypomethylated CpGs were associated with regions that lost the H3K36me2 mark in NSD1-KO cells, while only 10.7% of the hypomethylated CpGs were associated with gained H3K36me2 (Supplementary Figure 2D). It should also be noted that 54% of the hypomethylated CpGs were associated with regions that did not change their H3K36me2, indicating that NSD1 may regulate DNA methylation by a different mechanism in addition to its enzymatic activity.

**Figure 2:**
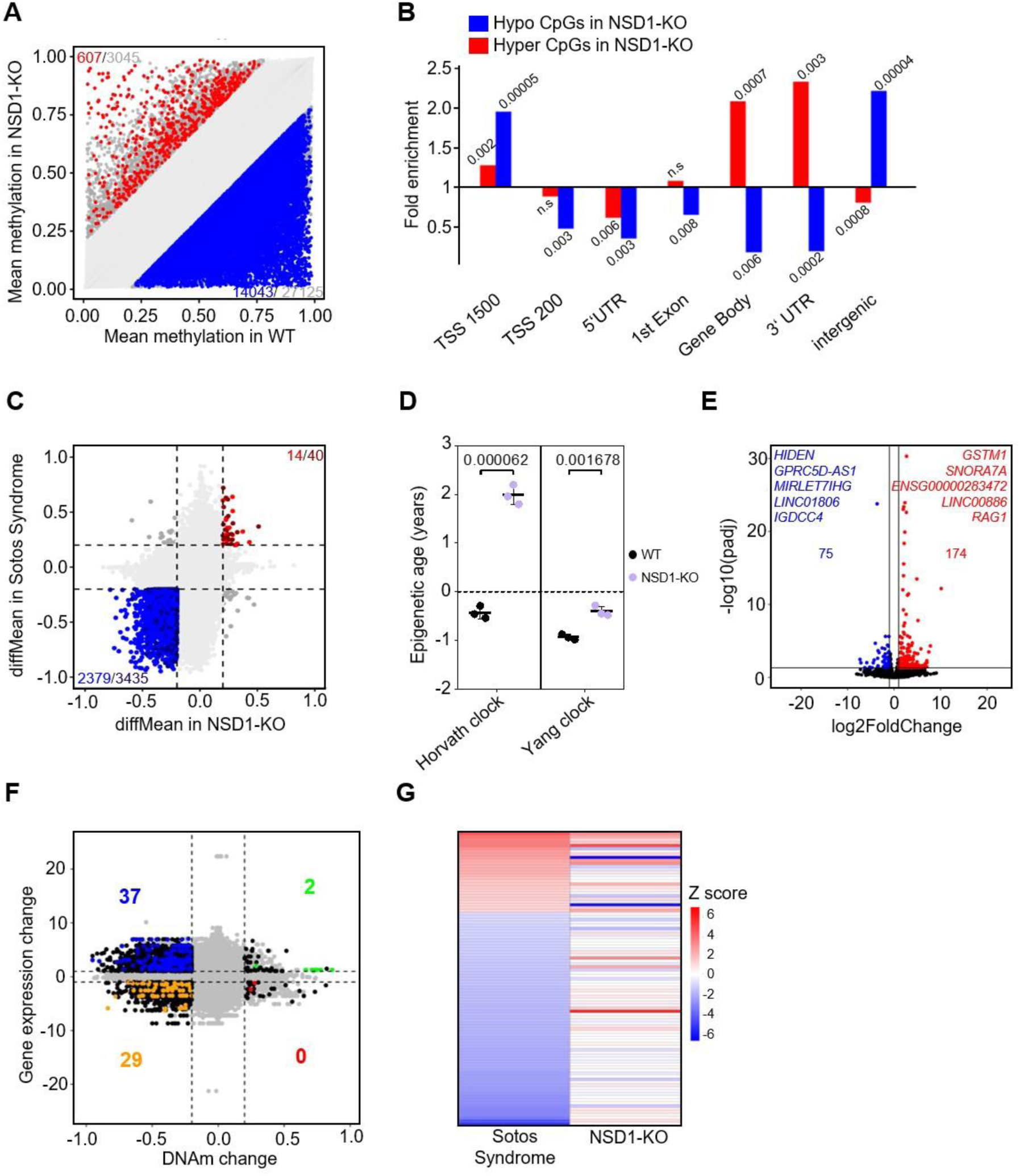
NSD1-KO induces global DNA hypomethylation and modest changes in transcription. **(A)** Scatter plot showing DNA methylation changes between NSD1-KO (n=3) and WT (n=3) with 14,043 hypomethylated sites and 607 hypermethylated sites (difference in mean methylation ≥20%, p-value≤0.05). **(B)** Association of hyper- and hypomethylated CpG sites in NSD1-KO iPSCs with genomic regions. Hypergeometric tests were used to calculate the significance, and p-values are depicted. **(C)** Comparison of DNA methylation in NSD1-KO iPSCs and Sotos syndrome samples. Mean Δbeta values of significantly different CpGs (difference in mean methylation ≥20%, p-value≤0.05) between NSD1-KO and WT are plotted against mean Δbeta values of significantly different CpGs (difference in mean methylation ≥20%, p-value≤0.05) between Sotos sydrome and healthy donors. Overlapping hypomethylated (blue) and hypermethylated (red) CpGs are indicated.**(D)** Epigenetic age calculation in WT and NSD1-KO iPSCs using the Yang clock and the Horvath clock. Age is depicted in years. Significance was determined by a paired t-test, and p-values are depicted. **(E)** Volcano plot of RNA-Sequencing analysis of NSD1-KO (n=3) and WT (n=3) iPSCs. 174 genes were significantly upregulated and 75 genes were significantly downregulated (>2-fold change, adj. p-value<0.05). The five genes exhibiting the most pronounced upregulation or downregulation are highlighted. **(F)** Association between DNA methylation levels and corresponding gene expression in WT and NSD1-KO iPSCs. Each data point represents a gene-CpG pair, with potential duplication of genes and CpGs. **(G)** Correlation of differentially expressed genes in NSD1-KO iPSCs and blood samples from patients with Sotos syndrome. The heat map represents normalized expression of differentially expressed genes in Sotos syndrome [18] in Sotos syndrome samples and NSD1-KO samples.

The overgrowth disorder Sotos syndrome is also characterized by a genome-wide loss of DNA methylation [21]. Brennan et al. identified the hypomethylation at cg07600533 as a diagnostic marker for Sotos syndrome [29], and our results showed significant hypomethylation at cg07600533 (Supplementary Figure 2E). This prompted us to compare global DNA methylation data in Sotos syndrome and NSD1-KO iPSCs. We used the DNA methylation data from blood of healthy donors and Sotos syndrome patients ([29], GSE191276) and observed that despite arising from different source material (iPSCs *versus* blood), 69.25% of CpGs hypomethylated in Sotos syndrome were also hypomethylated in NSD1-KO cells (Figure 2C), indicating that our NSD1-KO cells may recapitulate Sotos syndrome phenotypes. Previous reports have suggested that growth disorders such as TBRS and Sotos syndrome are associated with accelerated epigenetic age [22]. To determine whether a similar trend is observed in NSD1-KO cells, we computed epigenetic age using the Yang clock and the Horvath clock [30,31]. We observed a modest but significant epigenetic age acceleration in NSD1-KO iPSCs compared to WT cells (Figure 2D).In addition to overgrowth disorders, inactivating mutations in NSD1 have also been associated with cancers such as head and neck squamous cell carcinoma (HNSC) and lung squamous cell carcinoma (LUSC) [32], both of which have been characterized by global DNA hypomethylation. To determine whether similar overlaps as with Sotos syndrome samples could be detected in the cancer samples, we compared the DNA methylation profiles of HNSC and LUSC samples [32] with NSD1-KO. However, we did not find overlaps between the CpGs hypomethylated in either of the cancer samples and NSD1-KO cells (Supplementary Figure 2F, G).

RNA sequencing analysis of WT and NSD1-KO iPSCs revealed differences in the transcriptome based on the principal component analysis (PCA) (Supplementary Figure 2H). However, despite the substantial number of differentially methylated CpG sites identified, only a small number of genes showed significantly altered expression in NSD1-KO iPSCs. 174 genes were significantly upregulated, with *GSTM1*, *SNORA7A*, *ENSG00000283472*, *LINC00886*, and *RAG1* representing the five most highly upregulated genes. Among the 75 significantly downregulated genes, long noncoding RNAs (lncRNAs) exhibited the most substantial decrease, with *HIDEN, GPRC5D-AS, MIRLET7IHG, LINC01806* and *IGDCC4*, being the top 5 downregulated genes. (Figure 2E). Since NSD1 has a catalytically independent function as a transcriptional coactivator [15], we combined DNA methylation with transcriptomic data to better understand the regulation of the transcriptome through NSD1-mediated H3K36me2 and subsequent DNA methylation changes. Hypermethylated CpG sites overlapped just with 2 upregulated genes, while the hypomethylated CpGs overlapped with both upregulated (37) and downregulated (29) genes (Figure 2F), indicating a separate function of NSD1 in the regulation of gene expression and DNA methylation.

The DNA methylation profile of NSD1-KO cells partially recapitulated that of Sotos syndrome; however, comparative analysis revealed little overlap between genes dysregulated in Sotos syndrome and those in NSD1-KO iPSCs (Figure 2G). This results indicate that NSD1 may play distinct roles in regulating DNA methylation and transcription. Our data also suggest that, the DNA methylation profiles, but not the transcription profiles associated with developmental disorders, may already be partially established early in development. This recapitulation is not observed for NSD1-deficient cancer samples.

### NSD1-KO iPSCs exhibit endodermal and mesodermal differentiation defects

Following the detailed characterization of NSD1-KO iPSCs, their differentiation potential towards the three germ layers, endoderm, mesoderm and ectoderm was investigated. In an undifferentiated state, NSD1-KO iPSCs persisted in an iPSC-like morphology with unaffected expression of the pluripotency marker *OCT4* (Supplementary Figure 3A). Expression of germ layer-specific markers, *GATA6* (endoderm), *TBXT* (mesoderm), and *PAX6* (ectoderm) was evaluated in qRT-PCR analysis (Figure 3A). Upon endodermal differentiation, *GATA6* expression was significantly reduced in NSD1-KO cells compared to the WT, indicating an endodermal differentiation defect. *TBXT* and *PAX6* expression was not significantly different upon mesodermal and ectodermal differentiation, respectively. This trend was corroborated by immunophenotypic analysis, where GATA6 expression was almost absent in NSD1-KO cells while most of the WT cells were GATA6-positive upon endodermal differentiation. No considerable differences in protein expression of WT *versus* NSD1-KO iPSCs were observed in ectoderm with the mesodermal marker TBXT appearing increased in NSD1-KO cells (Figure 3B), Notably, the loss of H3K36me2 in endodermal marker genes, especially in intronic and intergenic regions, was consistent with the impaired differentiation of NSD1-KO iPSCs (Supplementary Figure 3B).

**Figure 3:**
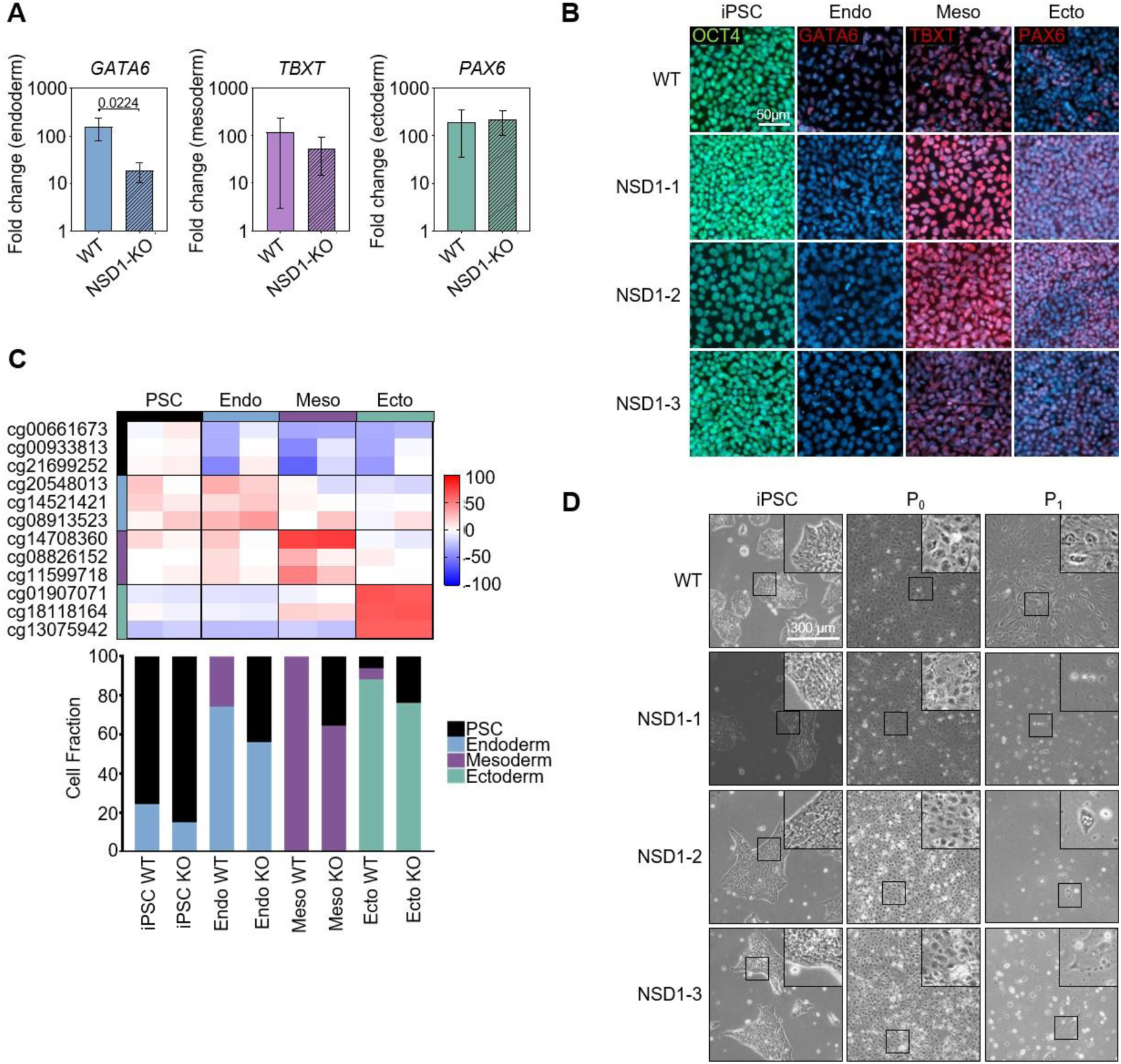
Endodermal and mesodermal differentiation capacity is impaired in NSD1-KO iPSCs. **(A)** qRT-PCR of germ layer-specific markers in endoderm (*GATA6*), mesoderm (*TBXT*), and ectoderm (*PAX6*). The trilineage differentiation was performed in triplicate for WT and the three NSD1-KO clones. Statistical analysis was performed with an unpaired t-test. P-value is depicted. **(B)** Immunofluorescence analysis of differentiation into the three germ layers. Antibodies targeting OCT4 (iPSCs), GATA6 (endoderm), TBXT (mesoderm), and PAX6 (ectoderm) were used. **(C)** Differentiation scores of selected CpG sites to evaluate the germ-layer specific differentiation potential of WT and NSD1-KO iPSCs. Each square represents the DNA methylation difference at a germ-layer-specific CpG site relative to reference values of PSCs. Below, the deconvolution shows estimates of the lineage-specific cell fractions during differentiation calculated as shown in [33].**(D)** Phase contrast microscopy images of WT and NSD1-KO iPSCs during iMSC differentiation.

We further validated the differentiation defects with DNA methylation data using the “PluripotencyScreen” [33]. Methylation values of 12 CpG sites – 3 CpGs per germ layer and pluripotent cells – provide a basis for deconvolution analyses to estimate the proportions of cells differentiating into each germ layer. The DNA methylation pattern in NSD1-KO iPSCs resembled an undifferentiated state, similar to WT iPSCs (Figure 3C). Following directed differentiation, the DNA methylation signature of NSD1-KO cells diverged markedly from WT cells, with a high fraction of differentiated cells still retaining pluripotency signatures, indicating impaired differentiation, particularly towards endodermal and mesodermal lineages (Figure 3C). When focusing only on pluripotency-associated CpG sites, NSD1-KO cells across all differentiation directions exhibit a DNA methylation pattern closer to a pluripotent state, compared to WT cells, as illustrated by the Pluripotency Score (Supplementary Figure 3C). Interestingly, differentiation of NSD1-KO iPSCs towards induced mesenchymal stromal cells (iMSCs) was not possible, reinforcing a role of NSD1 in late mesodermal differentiation (Figure 3D). Taken together, our results indicate that NSD1-mediated H3K36 dimethylation is an essential factor for early germ layer differentiation, especially in the endodermal and mesodermal lineages.

### NSD1 regulates endodermal differentiation via the HIDEN lncRNA

It has been previously reported that the loss of NSD1 leads to mesodermal defects [14]. We therefore decided to further investigate the role of NSD1 in endodermal differentiation. While NSD1-KO leads to modest changes in transcription, one of the most downregulated genes was the long non-coding RNA (lncRNA) *HIDEN* (ENSG00000253507; Transcript ID ENSG00000253507.5). Lu *et al*. reported that *HIDEN* is important for mRNA stability of the WNT receptor FZD5, positioning it as a crucial modulator for endodermal differentiation. Notably, knockout of *HIDEN* resulted in impaired endodermal differentiation capacity of human ESCs [27].

We confirmed the reduction of *HIDEN* expression in NSD1-KO by qRT-PCR and observed significantly reduced expression compared to WT cells in the pluripotent state and after endodermal differentiation (Figure 4A). Further investigating the mechanisms of NSD1-mediated regulation of *HIDEN* expression, we found that the *HIDEN* region exhibited an enrichment in H3K36me2 in WT iPSCs, which was significantly diminished in NSD1-KO cells already in the pluripotent state, especially in clones 2 and 3 (Figure 4B). CpG sites surrounding *HIDEN* promoter region and gene body (cg25940523, cg21410491, cg01065516, cg26980111, cg02642958), however, did not show a significant change in DNA methylation levels (Figure 4C). We investigated whether HIDEN KO definitive endoderm cells [27] and NSD1-KO cells would exhibit similar gene expression profiles by comparing RNA-Seq data and observed that genes up- or down regulated in HIDEN-KO were also similarly up-or down regulated in NSD1-KO (Figure 4D), reinforcing the role of HIDEN in NSD1-mediated endodermal differentiation.

**Figure 4:**
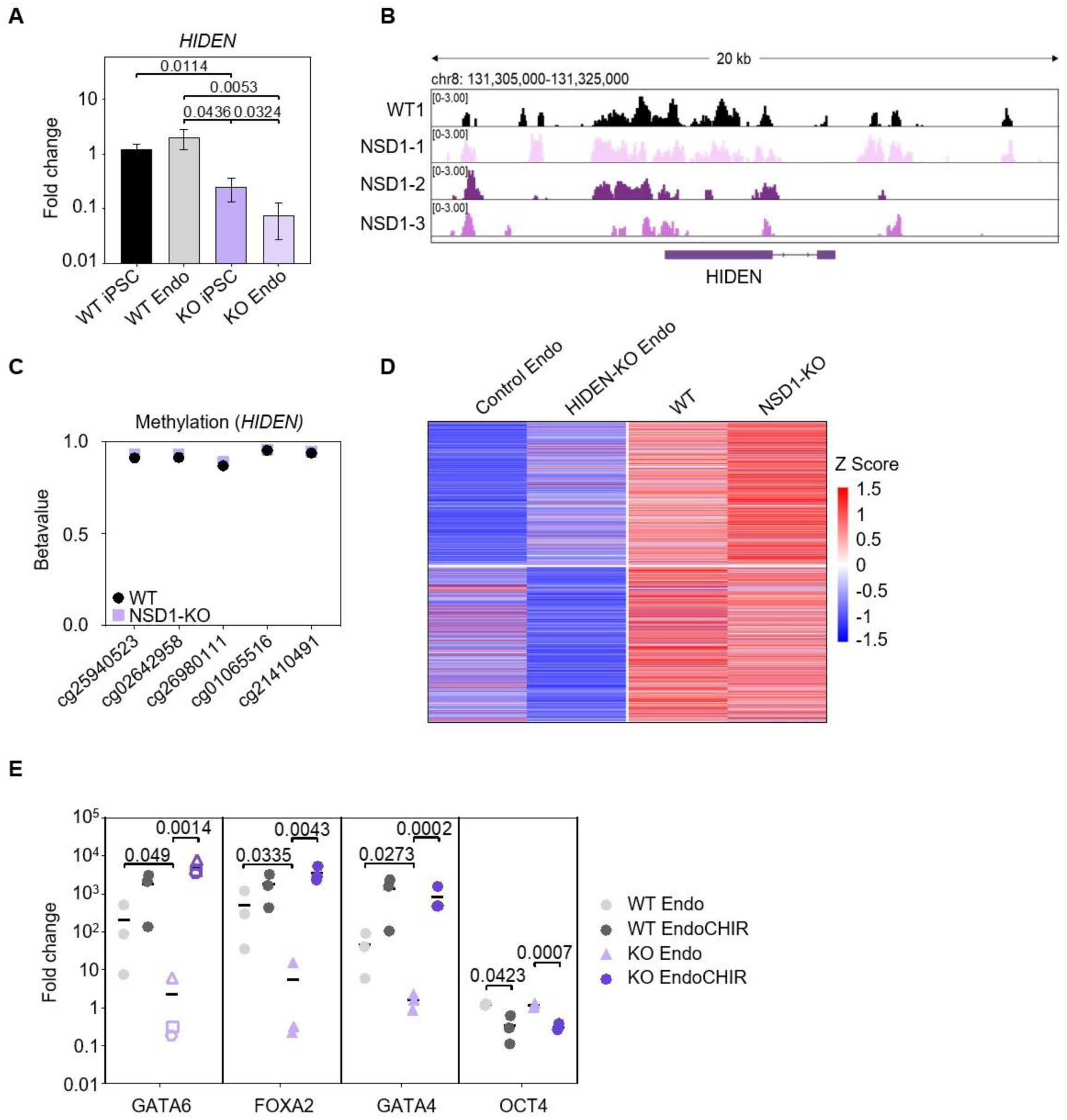
NSD1-mediated early differentiation is induced via lncRNA HIDEN. **(A)** Gene expression of *HIDEN* in undifferentiated WT and NSD1-KO iPSCs and after endodermal differentiation. Fold changes were normalized to *GAPDH*. Statistical analysis was performed with an unpaired t-test. P-values are depicted. **(B)** Visual representation of H3K36me2 enrichment at the *HIDEN* genomic region for WT and NSD1-KO iPSCs. **(C)** Beta-values of CpG sites close to lncRNA HIDEN in WT and NSD1-KO iPSCs. **(D)** Comparative analysis of differentially expressed genes in endodermally differentiated HIDEN-KO hESCs [27] and NSD1-KO iPSCs. **(E)** Expression analysis of endodermal markers *GATA6*, *FOXA2*, and *GATA4* and pluripotency marker *OCT4* in endodermal differentiated WT and NSD1-KO iPSCs without and with treatment with the WNT agonist CHIR. Fold change compared to undifferentiated iPSCs, after normalization with *GAPDH* is plotted. Statistical analysis was performed with an unpaired t-test. P-values are depicted.

Previous studies indicated that treatment with the WNT agonist CHIR was able to rescue the endodermal defect caused by HIDEN-KO [27]. Interestingly, we were also able to recover endodermal differentiation as measured by significant upregulation of *GATA6*, *FOXA2*, and *GATA4* upon treatment of NSD-1 KO cells with CHIR (Figure 4E). This is consistent with our hypothesis that NSD1-mediated H3K36 dimethylation regulates *HIDEN* expression and thereby regulates endodermal germ layer differentiation of human iPSCs.

## Discussion

Human iPSCs provide unparalleled potential for the modeling of diseases, especially developmental disorders [34]. We attempted to generate human iPSCs functionally deficient in NSD1, the major histone methyltransferase responsible for establishing the intergenic H3K36me2 modification, mutations in which lead to overgrowth disorders such as Sotos syndrome. While frameshift mutations were confirmed by Sanger sequencing, we did not see a complete loss of the H3K36me2 mark. While NSD1 is the predominant H3K36me2 transferase, studies have identified other NSD family proteins, such as NSD2 and NSD3, that establish this modification [35]. The activity of NSD2 and 3 may explain the residual H3K36me2 we observe in our cells. ChIP-Seq analysis revealed a significant reduction of the H3K36me2 mark in NSD1-KO cells, especially at enhancers. This is in line with previous studies that implicate NSD1 in enhancer function [16,25]. While NSD1-deficient mice and mouse ESC lines have been generated before [14–16,25,29], we report novel human iPSC lines with functionally deficient NSD1, which may be used as model systems to further study the acquisition of the developmental defects occurring in overgrowth disorders caused by NSD1 mutations.

As also seen in Sotos syndrome, NSD1-KO leads to severe DNA hypomethylation but only a modest change in transcription [18], with more than half the CpGs hypomethylated in Sotos syndrome also hypomethylated in NSD1-KO iPSCs. There was little overlap between the differentially expressed genes, however, indicating that the DNA methylation profiles resulting from a loss of NSD1 may already be partially established early in development, and the gene expression patterns may be differentiation-specific. Our study revealed a small but significant increase in epigenetic age, in contrast to previous reports that indicate that overgrowth disorders are associated with a severe acceleration of epigenetic age [22]. While it should be noted that iPSCs are epigenetically rejuvenated and may not accurately reflect an acceleration of epigenetic age, the perceived acceleration in overgrowth disorders may be a readout of global hypomethylation, and functional studies may determine whether the overgrowth phenotype corresponds to functional aging.

One of the key findings of our study is that NSD1 KO iPSCs retain pluripotency markers. In fact, we see DNA methylation levels similar to pluripotent cells even after differentiation in the three germ layers, resulting in a higher pluripotency score. A previous report has suggested that H3K36me is a barrier to reprogramming, and suppression of H3K36 methylation enhanced the reprogramming of mouse embryonic fibroblasts (MEFs) associated with a PRC2-dependent silencing of mesenchymal genes and increased expression of epithelial and pluripotency genes [36,37]. We observe a similar push towards pluripotency upon loss of NSD1. Additionally, we observe defects in the differentiation potential in NSD1 KO cells, especially along the endoderm and mesoderm lineages. Differentiation of our NSD1-KO iPSCs into iMSCs was impaired, consistent with previously reported mesodermal defects in developing NSD1^-/-^ mice [14]. NSD1 KO in mouse ESCs exhibited differentiation defects in all three germ layers, and auxin-degron-based NSD1 degradation showed defects in cardiomyocyte differentiation and neural specification in mESCs [15,16]. Furthermore, NSD1 deficiency in mesenchymal progenitors lead to impaired chondrogenic differentiation and skeletal growth [38]. We further identify the endoderm-specific desert lncRNA *HIDEN* as being one of the most highly repressed gene upon NSD1 KO and demonstrate that NSD1 may regulate endodermal differentiation mediated by *HIDEN* and the Wnt signaling pathway. While H3K36me2 levels decrease upstream and in the intronic regions of *HIDEN*, the CpGs do not show a change in methylation. This indicates the NSD1 regulated *HIDEN* expression via mechanisms independent of DNA methylation and may rely on cross-talk between H3K36me2 and other histone modifications for regulating expression.

Integrating our results with previous reports that implicate NSD1 regulating cell fate determination through enhancer binding and regulation of bivalent gene expression [16,25,29], we can determine a dual mechanism of NSD1 function. NSD1 regulates transcription of developmental genes such as *HIDEN*, by precluding these regions from silencing and hence maintaining gene expression profiles, conducive to proper cell fate determination. NSD1 also maintains cell type specific DNA methylation profiles; global DNA hypomethylation occurs in the absence of NSD1, especially at intergenic regions, early in development, which may ultimately lead to lineage specific defects as seen in our cells and in Sotos syndrome. The NSD1-KO human iPSC lines therefore, may provide valuable insights into early differentiation defects that could elucidate the acquisition of lineage-specific phenotypes in overgrowth disorders.

## Methods

### Cell culture and directed differentiation

Three independent human induced pluripotent stem cell lines were reprogrammed from bone marrow-derived mesenchymal stromal cells by episomal plasmids – hPSCreg: UKAi009-A (wildtype (WT)1), UKAi010-A (WT2), and UKAi011-A (WT3). All samples were taken after informed and written consent using guidelines approved by the Ethics Committee for the Use of Human Subjects at the University of Aachen (permit number: EK128/09). All methods were performed in accordance with the relevant guidelines and regulations. Maintenance culture of WT iPSCs was performed on tissue culture plastic (TCP) coated with vitronectin (VTN, 0.5 µg/cm^2^; Stem Cell Technologies, Vancouver, Canada) or with Biolaminin 521 LN (LMN, 0.9 μg/cm2, BioLamina, Sundbyberg, Sweden) for NSD1-KO iPSCs. IPSCs were cultured in StemMACS iPS-Brew XF (Miltenyi Biotec GmbH, Bergisch Gladbach, Germany). For directed differentiation towards the three germ layers, endoderm, mesoderm, and ectoderm, the STEMDiff™ Trilineage Differentiation Kit (Stem Cell Technologies, Vancouver, Canada) was used according to the manufacturer’s instructions. Differentiation into endodermal and mesodermal direction lasted for 5 days and into ectodermal direction for 7 days. As an undifferentiated control, iPSCs were maintained in StemMACS iPS-Brew XF for 5 days. Endodermal rescue experiments were performed by treatment of cells with endoderm differentiation medium containing 2μM CHIR99021 (CHIR, Tocris). Differentiation towards induced mesenchymal stromal cells (iMSCs) was performed according to a protocol from [26]. The differentiation was initiated by switching from iPS-Brew XF to DMEM low glucose supplemented with 10% human platelet lysate, when a confluency of 60-70% was reached. Cells were maintained for 7 days (P0) and then passaged to 0.1% gelatine-coated TCP (P1). Since the NSD1-KO cells were not viable following passaging, the differentiation was not continued.

### CRISPR/Cas9 mediated NSD1 modification

For the generation of NSD1-KO cell lines, WT1 iPSCs were transfected with a CRISPR/Cas9 nuclease approach. The guide RNA was designed to target exon 3 within the human NSD1 gene (ENSG00000165671, gRNA sequence: AATTCAAGAGACGCCCATGG). The ribonucleoprotein (RNP) was assembled using Alt-R CRISPR/-Cas9 crRNA, Alt-R CRISPR/-Cas9 tracrRNA, and Alt-R HiFi Sp. Cas9 Nuclease (all IDT, Coralville, USA), and transfected into WT1 iPSCs using the NEON transfection system (1300 V, 30 ms pulse width, 1 pulse; Thermo Fischer Scientific, Waltham, USA). Following the electroporation, transfected cells were seeded on a coating of Biolaminin 521 LN in StemMACS iPS-Brew XF, supplemented with Rho-associated protein kinase (ROCK) inhibitor Y-27632 (10 µM, Abcam, Cambridge, UK) and 1x CloneR (Stem Cell Technologies, Vancouver, Canada). Four days later, cells were singularized and seeded on mouse embryonic fibroblasts (MEFs). After eight days, individual iPSC colonies were picked and seeded on MEFs for an additional passage and subsequently expanded in a feeder-free environment on vitronectin-coated plates. Expanded clones were screened for mutations in exon 3 via Sanger Sequencing (Eurofins Genomics, Ebersberg, Germany; sequencing primer: AAAGTCTACGCCACTGAAGTATG). The culture of the three NSD1-KO clones was changed to LMN coating, after a significant increase in the viability of NSD1-KO iPSCs on LMN was seen.

### Growth curves

Equal numbers of cells in single-cell suspensions (150,000 cells; WT1, NSD1-KO clone 1, 2, 3) were seeded onto TCP coated with either VTN or LMN. For cell counting, cultures were dissociated using Accutase (Stem Cell Technologies, Vancouver, Canada), collected by centrifugation, and counted using the Countess 3 automated cell counter (Thermo Fischer Scientific, Waltham, USA). Cell counts were determined every 24 hours over 4 days in triplicates.

### Western Blot

Total cell extracts from iPSCs were prepared in cold lysis buffer (RIPA buffer, 50 mM NaF, 1 mM Na_3_VO_4_, and 7× complete mini protease inhibitor). The protein concentration was determined by Bradford protein assay, measured in a photometer (Infinite 200 PRO, Tecan Trading AG, Switzerland). Of each sample, 15 µg protein was incubated in 4x SDS Protein Sample Buffer (50 mM Tris/HCl pH 6.8, 2% SDS, 0.01% Bromophenol blue, 2.5% β-mercaptoethanol and 10% glycerol) for 5 min at 99°C. The samples were separated in 12% Mini-PROTEAN TGX Precast Protein Gels (Bio-Rad, München, Germany) and transferred to a polyvinylidene fluoride membrane (PVDF; Merck Millipore, Burlington, USA). The membrane was blocked in 4%w/v bovine serum albumin (BSA) for one hour and then incubated in primary antibodies against H3K36me2 and GAPDH overnight at 4°C (All antibodies indicated in Supplementary Table 1). Incubation with secondary antibodies was done for 1h at room temperature (RT), and protein bands were visualized using a ChemiDoc XRS+ (Bio-Rad Laboratories, Hercules, USA).

### Immunophenotypic analysis

Fixation of the cells was performed with 4% paraformaldehyde for 20 min. Fixed cells were treated with PBS containing 1%w/v BSA and 0.1%v/v Triton-X-100 (Bio-Rad, München, Germany) for 1h at 4°C, and then incubated in primary antibodies (Supplementary Table 1) overnight at 4°C. Secondary antibody staining was performed at RT for 1h, and thereafter, nuclei were counterstained with Hoechst 33342. Fluorescence microscopy was performed with a Zeiss Axio Observer (Carl Zeiss, Oberkochen, Germany) using 20x or 40x or 63x objectives.

### Semi-quantitative reverse-transcriptase PCR

Total RNA was extracted with the NucleoSpin RNA Plus Kit (Macherey-Nagel, Düren, Germany) and quantified with a NanoDrop 2000 spectrophotometer (Thermo Scientific, Waltham, USA). Isolated RNA was converted into complementary DNA (cDNA) using the High-Capacity cDNA Reverse Transcription Kit (Applied Biosystems, Waltham, USA). To analyze the gene expression of target genes, a semi-quantitative real-time PCR (qRT-PCR) was performed, using TaqMan gene expression master mix and gene-specific TaqMan assays (Supplementary Table 2). The qRT-PCR was performed in a StepOnePlus Real-Time PCR cycler. For analysis of aberrant endodermal differentiation, qRT-PCR was carried out using iTaq Universal SYBR Green Supermix (Bio-Rad, München, Germany) and primers mentioned in Supplemental Table 2 in a CFX Duet Real-Time PCR System (Bio-Rad, München, Germany). All reactions were pipetted in technical duplicates. After performing the qRT-PCR, results were analyzed with the ΔΔCt method, normalizing to the housekeeping gene *GAPDH*.

### Transcriptomic analysis

Total RNA from iPSCs was isolated with the NucleoSpin RNA Plus Kit. For each sample, 700 ng RNA were sent to Life&Brain GmbH (Bonn, Germany), where library preparation (QuantSeq 3′-mRNA) and RNA-sequencing were performed on a NovaSeq6000 sequencer. Quality of FASTQ files was quantified using FastQC, and adaptor sequences and low-quality reads were trimmed using Trim Galore!. Alignment of the reads was done using STAR (hg38 genome build), and transcript quantification was performed using Salmon. The resulting count matrices were normalized by size factor and dispersion using the DESeq2 package in R. Differential gene expression analysis between NSD1-KO and WT samples was performed using Negative Binomial GLM fitting and Wald-test, and p-values were adjusted with the Benjamini-Hochberg procedure. Genes with adjusted p-value <0.05 and absolute log2fold change >1 were considered as differentially expressed. Graphical representation of gene expression differences was done using the Z-Score, computed from normalized counts according to the formula: (counts – mean(counts))/sd(counts). Lists of signature genes for each germ layer (ectoderm, endoderm and mesoderm) were obtained from the literature [33]. A list of signature genes for Sotos Syndrome was obtained from [18].

### Pyrosequencing

Genomic DNA (gDNA) was isolated from differentiated and undifferentiated iPSCs using the NucleoSpin Tissue Kit (Macherey-Nagel, Düren, Germany). 500 ng gDNA were bisulfite converted with the EZ DNA methylation Kit (Zymo Research), and target sequences were PCR amplified using the Pyromark PCR Kit (Qiagen) with 2.5 mM Mg^2+^ and a primer concentration of 0.3 µM (Supplemental Table 2). Pyrosequencing was performed on a Q48 pyrosequencer (Qiagen) and analyzed using the Pyromark Q48 autoprep software (Qiagen).

### DNA methylation profiling

Genomic DNA was bisulfite converted and analyzed with the Illumina human EPIC methylation microarray version 2 (EPIC v2) for three WT and three NSD1-KO clones (all analyzed at Life and Brain GmbH, Bonn, Germany). For comparison of NSD1-KOwith Sotos syndrome, LUSC and HNSC, public datasets from [29,32] were used. Initial quality control of the DNA methylation data was carried out using the minfi package (v1.50.0) [39]. The preprocessing, including dye bias correction, quality mask filtering, NOOB normalization, and detection p-value calculation, was performed using the SeSAMe package (v1.22.2) [40]. We excluded CpG probes that failed in 10% or more of the samples, non-CpG probes and probes on the X and Y chromosomes. The data was converted into a GenomicRatioSet for further analysis with minfi-based functions. For the multidimensional scaling (MDS) plots and differential methylation analysis, we used Limma (v3.60.0) [41], considering probes with a mean beta value difference of ≥ 0.2 and adjusted p-values (Benjamini-Hochberg) of ≤ 0.05 as significant. Heatmap was generated using the ComplexHeatmap package (v2.20.0), employing Pearson correlation as the distance measure and the “ward.D2” clustering method. Gene ontology analysis was conducted with the missMethyl R package (v1.38.0) [42]. The R packages: watermelon, ggplot2, ggrepel, ggbeeswarm, reshape2, ggExtra, gprofiler2, and ComplexHeatmap were used for graphical presentation. Epigenetic age was calculated using the Horvath multi-tissue clock [30] or the ‘Yang clock’ reported in [31].

### Chromatin immunoprecipitation sequencing

iPSCs were fixed with 1% formaldehyde and sonicated using the Covaris M200 (75W, 10%, 200 cycles/burst) for 30 minutes to obtain chromatin fractions ranging from 200 bp-1kbp. Sonication was performed by the Genomics Facility, IZKF, University Hospital, Aachen. The ChIP protocol was carried out according to the manufacturer’s recommendations (Active motif #53042). Library preparation and ChIP sequencing were carried out by Cologne Centre for Genomics. Fastq files were analyzed using the nextflow pipeline (nf-core/chipseq version 2.1.0). BigWig files, for each ChIP sample normalized to its input, were created from sortedBam files generated by nextflow (SAMtools), using bamCompare from deepTools (binSize 50). BigWig tracks were visualized using IGV [43,44]. Heatmaps and line plots were generated from the normalized bigWig files with computeMatrix from deepTools. GeneHancer BED GRCh38 file was obtained from GeneCards.org [45]. Downstream analysis of broadPeak files from nextflow (MACS3) was carried out in R using rtracklayer, GenomicRanges for overlapping ChIP-Seq with DNA methylation and RNA-Seq data, ChIPseeker for peaks annotation, and clusterProfiler for GO enrichment analysis of peaks.

## Author Contributions

A.H. generated and characterized the NSD1-KO iPSC lines under the supervision of K.Z.. A.H. and K.Z. performed experiments and generated samples for RNA sequencing and DNA methylation analysis. M.V.B. and E.D.T. performed bioinformatics analysis. M.M. generated samples for ChIP-Seq and performed analysis. D.P. designed the study, and D.P., K.Z., and W.W. supervised the experiments. A.H., K.Z. and D.P. wrote the manuscript, and all authors approved the final version.

