## Supplemental material for "NSD1 governs H3K36me2-mediated DNA methylation and drives differentiation of human iPSCs by regulating ‘HIDEN’ lncRNA expression"

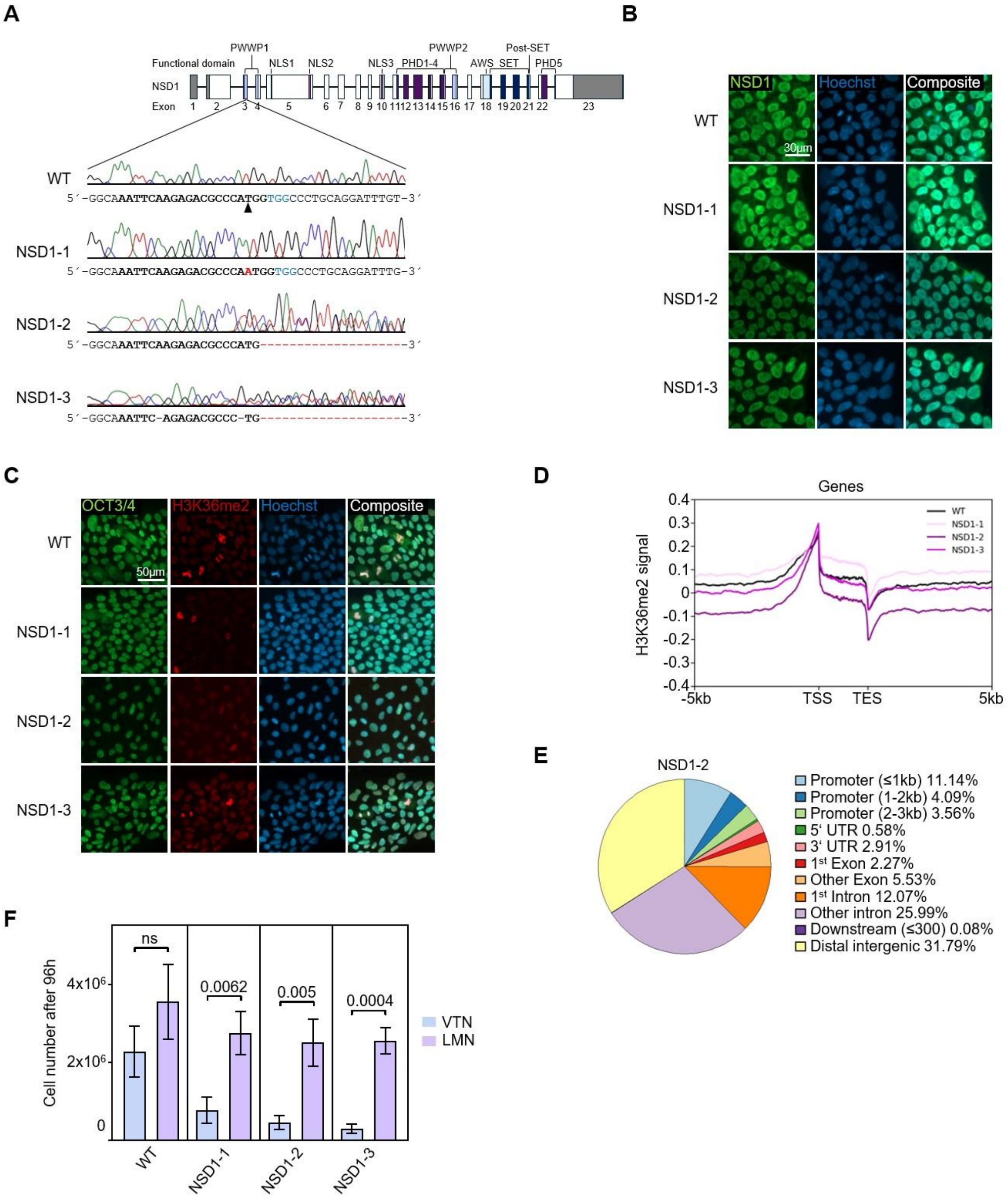

### Supplementary Figure 1: Generation of NSD1-KO iPSCs

**(A)** Domain structure of human *NSD1* and CRISPR/Cas9 strategy targeting exon three of canonical *NSD1*. Three iPS cell lines (NSD1-1, NSD1-2, NSD1-3) with homozygous or heterozygous frameshift mutations were generated. **(B)** Immunophenotypic analysis of WT and NSD1-KO iPSCs. Cells were stained with an NSD1 antibody and nuclei were counterstained with Hoechst. **(C)** Immunophenotypic analysis of WT and NSD1-KO iPSCs. Cells were stained with an OCT3/4 and H3K36me2 antibody and nuclei were counterstained with Hoechst. (**D)** Line plot showing H3K36me2 ChIP-seq signal over transcription start site (TSS) and transcription end site (TES) ±5 kbp for WT and NSD1-KO clones. (**E)** Pie chart indicating gene features of regions that lose H3K36me2 in NSD1-KO compared to WT. The values for the exemplary clone 2 are depicted. **(F)** Quantification of the total cell number of WT and NSD1-KO iPSCs growing on vitronectin or laminin after 96h. Statistical analysis was performed using an unpaired t-test, and p-values are depicted.

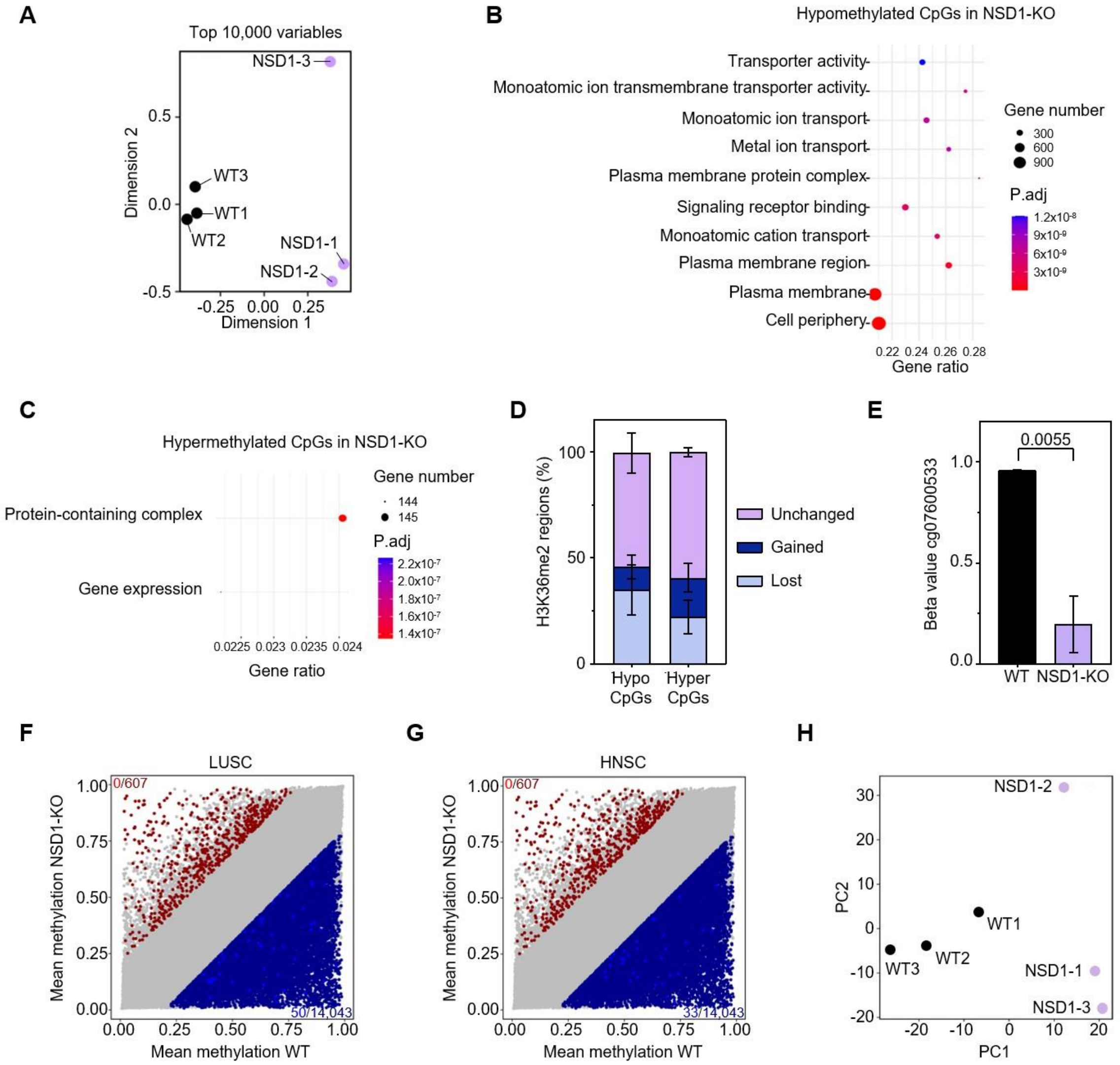


**Supplementary Figure 2: DNA methylation and gene expression analysis of NSD1-KO cells**

**(A)** MDS plot of the top 10,000 variables of DNA methylation data from three WT and three NSD1-KO iPSC clones. **(B)** Gene set enrichment analysis of hypomethylated CpG sites in NSD1-KO indicates significant enrichment of genes associated with the plasma membrane and ion transport. **(C)** Gene set enrichment analysis of hypermethylated CpG sites in NSD1-KO. **(D)** Classification of CpGs on H3K36me2 changes. Hyper or Hypomenthylated CpGs in NSD1-KO are classified based on gain, loss or no change in H3K36me2 modification in NSD1-KO. **(E)** Bar plot showing the DNA methylation of the Sotos syndrome diagnostic CpG site in WT and NSD1-KO iPSCs. Statistical analysis was performed using an unpaired t-test, and p-value is depicted. **(F-G)** Comparison between differentially methylated CpGs in NSD1-KO iPSCs and LUSC (F) and HNSC (G). Only 50 and 33 CpGs hypomethylated in NSD1-KO were also hypomethylated in LUSC and HNSC respectively (light blue). No overlaps were detected between the hypermethylated CpGs.**(H)** PCA of RNA sequencing results of three WT and three NSD1-KO iPSC clones.

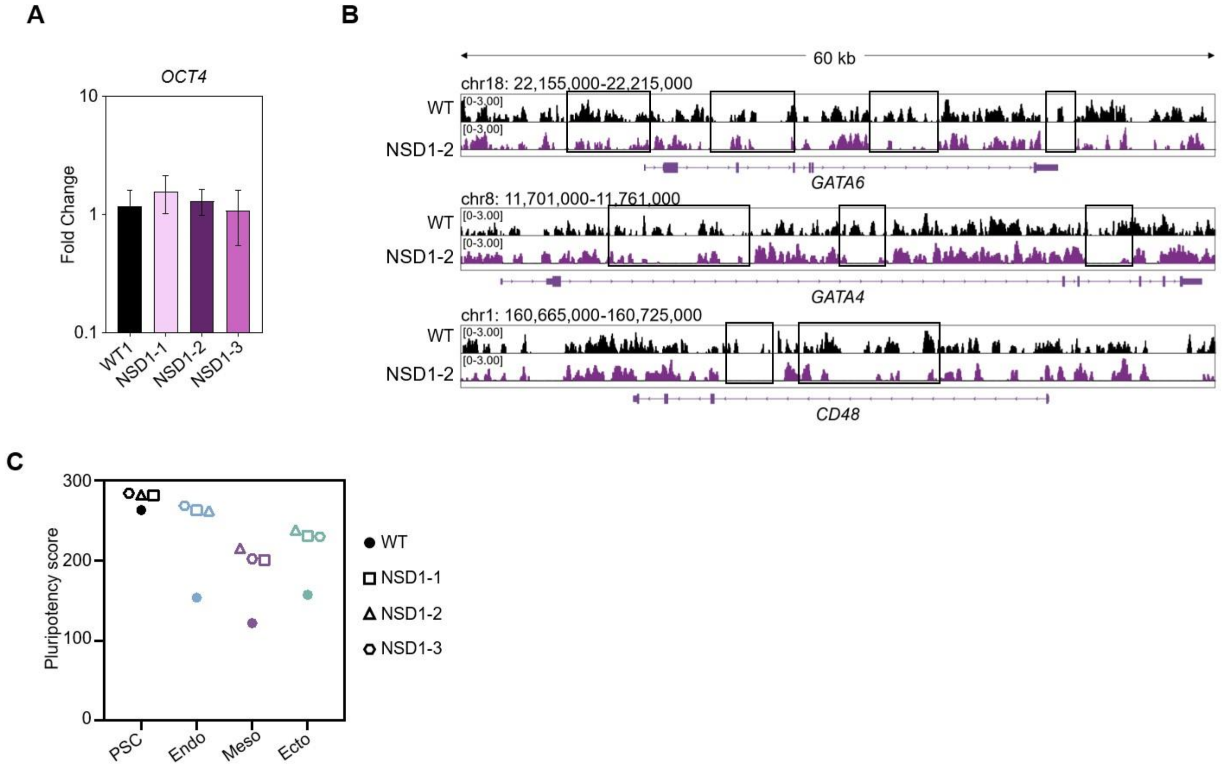


**Supplementary Figure 3: Differentiation defects in NSD1-KO**
**(A)** Gene expression of *OCT4* in undifferentiated NSD1-KO iPSCs. Fold changes were normalized to *GAPDH* and expression of *OCT4* in WT iPSCs. Statistical analysis was performed with an unpaired t-test.**(B)** Visual representation of H3K36me2 changes on select endodermal genes between WT and exemplary clone 2 for NSD1-KO.Representative regions that show loss or decrease in H3K36me2 in NSD1-KO are marked. (**C)** Pluripotency Score of WT and NSD1-KO iPSCs in undifferentiated state and upon endodermal, mesodermal, and ectodermal differentiation. The Pluripotency Score was calculated using the sum of the beta-values of three pluripotency-specific CpG sites (Schmidt *et al.*, 2023).

### Supplementary Table 1: Antibodies used in this study

| Primary antibodies | | | | | | |
|---|---|---|---|---|---|---|
| **Name** | **Host** | **Application** | **Dilution** | **Company** | **Clonality** | **Catalogue-Nr.** |
| Anti-H3K36me2 | Rabbit | WB | 1:2,500 | Abcam | polyclonal | ab9049 |
| Anti-H3K36me2 | Rabbit | ChIP | 4µg | Abcam | polyclonal | ab9049 |
| Anti-H3K36me2 | Rabbit | IF | 1:2,000 | Abcam | polyclonal | ab9049 |
| Anti-NSD1 | Rabbit | IF | 1:300 | Thermo Fisher Scientific | polyclonal | PA5-50857 |
| Anti-OCT3/4 | Mouse | IF | 1:50 | Santa Cruz | monoclonal | sc-5279 |
| Anti-GATA6 | Rabbit | IF | 1:1,600 | Cell Signaling | monoclonal | 5851S |
| Anti-Brachyury | Goat | IF | 1:100 | R&D systems | polyclonal | AF2085 |
| Anti-PAX6 | Mouse | IF | 1:100 | Santa Cruz | monoclonal | sc-53108 |
| **Secondary antibodies** | | | | | | |
| **Name** | **Host** | **Target** | **Application** | **Dilution** | **Company** | **Catalogue-Nr.** |
| Alexa Fluor™ 594 | Rabbit | Goat | IF | 1:200 | Invitrogen | A-11012 |
| Alexa Fluor™ Plus 488 | Rabbit | Goat | IF | 1:200 | Invitrogen | A32731 |
| Alexa Fluor™ Plus 488 | Mouse | Goat | IF | 1:200 | Invitrogen | A32732 |
| Peroxidase AffiniPure™ | Rabbit | Goat | WB | 1:5,000 | Jackson ImmunoResearch | 111-035-003 |

### Supplementary Table 2: Primers used in this study

| | | |
|---|---|---|
| **Taqman assays** | | |
| **Target** | **Assay ID** | |
| *NSD1* | Hs00328315_m1 | |
| *POU5F1* | Hs04260367_gH | |
| *GATA6* | Hs00232018_m1 | |
| *T* | Hs00610080_m1 | |
| *PAX6* | Hs01088114_m1 | |
| *GAPDH* | Hs02758991_g1 | |
| **qRT-PCR Primers** | | |
| **Name** | **Sequence** | |
| *OCT4* Fwd | GGGGGTTCTATTTGGGAAGGTA | |
| *OCT4* Rev | ACCCACTTCTGCAGCAAGGG | |
| *GATA6* Fwd | CTCAGTTCCTACGCTTCGCAT | |
| *GATA6* Rev | GTCGAGGTCAGTGAACAGCA | |
| *FOXA2* Fwd | GCACTCGGCTTCCAGTATGCTG | |
| *FOXA2* Rev | TCACGGAGGAGTAGCCCTCG | |
| *GATA4* Fwd | GGCCTCTACATGAAGCTCCACG | |
| *GATA4* Rev | CTGAAGGAGCTGCTGGTGTCTT | |
| *TBXT* Fwd | CAGTGGCAGTCTCAGGTTAAGAAGGA | |
| *TBXT* Rev | CGCTACTGCAGGTGTGAGCAA | |
| *HIDEN* Fwd | TCACCGGTCCTCTTGTGTTG | |
| *HIDEN* Rev | TTCTTTTCCAAAGCCGCTGA | |
| *GAPDH* Fwd | GAAGTTGAAGGTCGGAGTC | |
| *GAPDH* Rev | GAAGATGGTGATGGGATTTC | |
| **Pyrosequencing primers** | | |
| **CpG site** | **Name** | **Sequence** |
| cg00661673 | cgSC1 Fwd | GGTTGGAGTGTATTGGTGTAA |
| | cgSC1 Rev | Biotin-AATCCCAACCTTTATACATATTAATTCTT |
| | cgSC1 Seq | GTTGAGATTATAGGTGTGA |
| cg00933813 | cgSC2 Fwd | AGGTTGGTTATGAATTTTTGGTTTTAAGTA |
| | cgSC2 Rev | Biotin- ATACCCTACCTTCCTTTCATTTATATTC |

| | | |
|---|---|---|
| | cgSC2 Seq | TTGGGATTATAGGTGTG |
| cg21699252 | cgSC3 Fwd | GATGTTGAGGGTTAGGGGGTAATT |
| | cgSC3 Rev | Biotin- CCTAAAACTCTAAAAATCTTTCTCCCTAAA |
| | cgSC3 Seq | TGAAGGTTTTTTTAGTTTTGA |
| cg20548013 | cgE1 Fwd | GAATAGTATATGGTTGGTTGGGAAAGT |
| | cgE1 Rev | Biotin- CCAAAAAAAAAAAATACCTTTACTATCACT |
| | cgE1 Seq | AGGAGTTATTTTATTATATTGGAG |
| cg14521421 | cgE2 Fwd | GGGATGTTGTGGATGGTAAAA |
| | cgE2 Rev | Biotin- ACTCCCACATCTAAACACCTAA |
| | cgE2 Seq | AGGGGTGTGGGAAGT |
| cg08913523 | cgE3 Fwd | GGGAGAGGGATTTATTATTAGGT |
| | cgE3 Rev | Biotin- ACCCCCTCCTTCAACTATAAT |
| | cgE3 Seq | GGTTTGAGAAAGAAGTTAG |
| cg14708360 | cgM1 Fwd | AGGGTAAGGTTGTTTTGTTTAGTTTAT |
| | cgM1 Rev | Biotin- TCATACCTTTAAAACCCACAACTAAAAT |
| | cgM1 Seq | ATTAGGGTTTTGGTTTTATT |
| cg08826152 | cgM2 Fwd | TGAGTTTGGTTAGTTTAGTTATAGGT |
| | cgM2 Rev | Biotin- CATCCCTAAAACAAACAAAAAACAATT |
| | cgM2 Seq | ATTTGTTGTTGAGGTTTTTAATA |
| cg11599718 | cgM3 Fwd | ATGGTTTGGTATAGAAAGTTTATGG |
| | cgM3 Rev | Biotin- ATACTTTCATCTCTTCTAATACCTTTAAC |
| | cgM3 Seq | GTTTTGTGGGTGGGG |
| cg01907071 | cgEC1 Fwd | GGGGTTTTGAAAGTAAATGTGT |
| | cgEC1 Rev | Biotin- TTCCAACTCACTAAAAAACACTTC |
| | cgEC1 Seq | AGTAAATGTGTTGAAAGTT |
| cg18118164 | cgEC2 Fwd | AGTGGGAGTAAATGAGTTTAGT |
| | cgEC2 Rev | Biotin- CAATTTCAAAATCTCCATCTCAAAATATCA |
| | cgEC2 Seq | TTTTAGGGTAAGAAAATATAGATAG |
| cg13075942 | cgEC3 Fwd | GGGAGATTTTAGTTTTTTTTGTAGGG |
| | cgEC3 Rev | Biotin- CCCAATATTATAATTCTTAACACCTCTCAT |
| | cgEC3 Seq | AGTTTTTTTTGTAGGGATTTT |
